# Dysfunctional vascular smooth muscle cells mediate early and late-stage neuroinflammation and Tau hyperphosphorylation

**DOI:** 10.1101/2021.04.13.439741

**Authors:** Jorge A. Aguilar-Pineda, Karin J. Vera-Lopez, Pallavi Shrivastava, Rita Nieto-Montesinos, Miguel A. Chávez-Fumagalli, Gonzalo Davila Del-Carpio, Badhin Gomez Valdez, Clint L. Miller, Rajeev Malhotra, Mark E. Lindsay, Christian L. Lino Cardenas

## Abstract

Despite the emerging evidence implying early vascular contributions to neurogenerative syndromes, the role of vascular smooth muscle cells (VSMCs) in the pathogenesis of Alzheimer’s disease is still not well understood. Herein, we show that VSMCs in brains of AD patients and the animal model of the disease, are deficient in multiple VSMC-contractile markers which correlated with Tau accumulation in brain arterioles. *Ex vivo* and *in vitro* experiments demonstrated that VSMCs undergo dramatic phenotypic transitions under AD-like conditions, adopting pro-inflammatory and synthetic phenotypes. Notably, these changes coincided with Tau hyperphosphorylation at residues Y18, T205 and S262. We also observed that loss of VSMC markers occurred in an age-dependent manner, and that expression of Sm22α and α-Sma proteins were inversely correlated with CD68 and Tau accumulation in brain arterioles of 3xTg-AD mice. Together, these findings further support the contribution of VSMCs in AD pathogenesis, and nominate VSMCs as potential novel therapeutic target in AD.

**Graphical Abstract:** 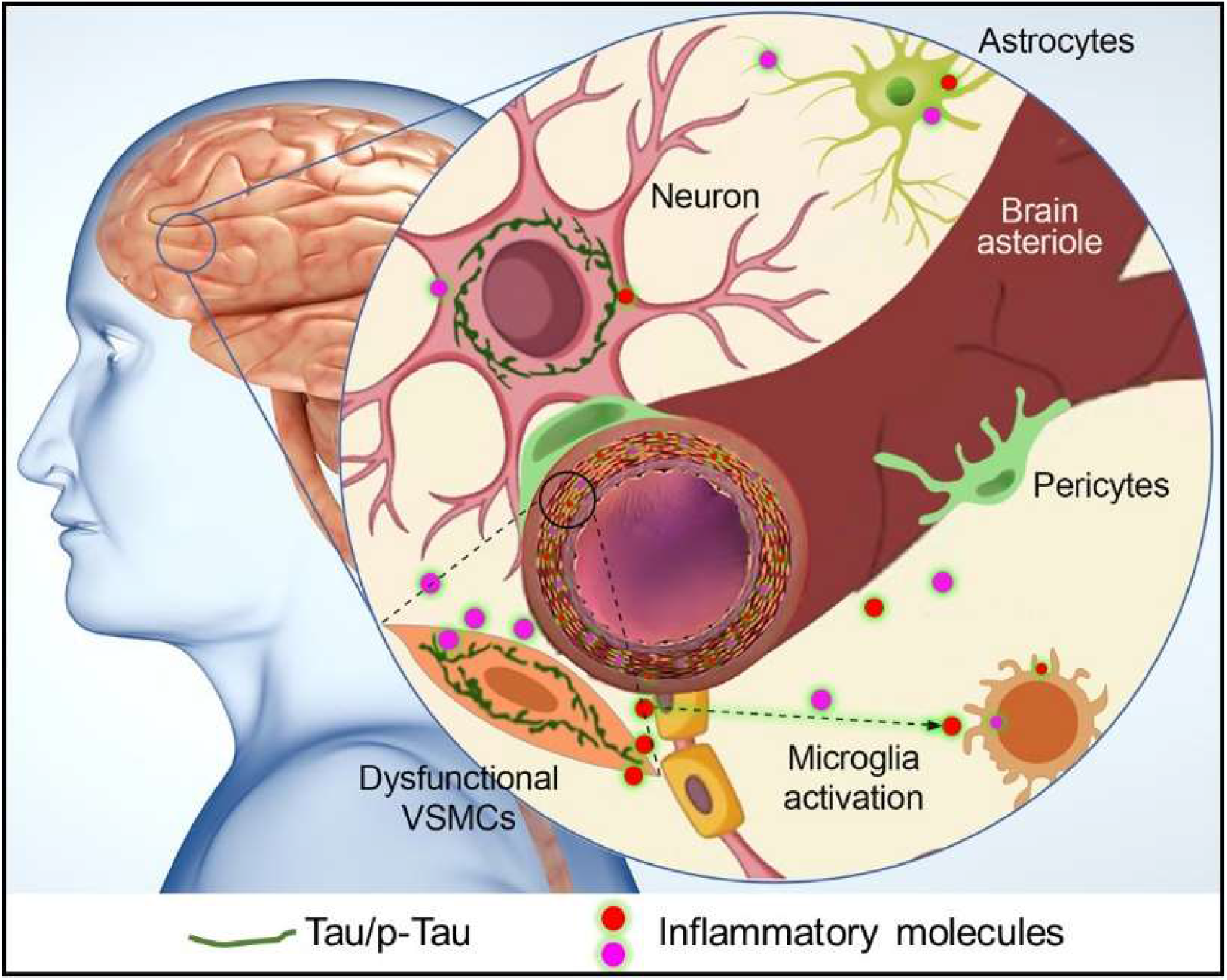

## INTRODUCTION

Alzheimer’s disease (AD) is a debilitating neurological disorder that leads to cognitive decline and dementia. The prevalence of AD patients is predicted to surge more than 150% in the next decade, in line with the increased aging in US population (>65 years old) (Morris et al., 2020; Alzheimer’s Association 2018). Tau neurofibrillary tangle (NFT) deposits are characteristic hallmarks of AD and it correlates closely with cognitive decline which begins to develop around two decades before clinical manifestations (Nirzhor et al., 2018; Leszek et al., 2013; Barthélemy et al., 2020). The microtubule-associated Tau protein (MATP) plays a key role in the morphogenesis and homeostasis of mature neurons. Also, multiple residues in the Tau protein (∼80aa) undergo post translational modifications (PTMs), which are critical for proper assembly and stability of the microtubule cytoskeleton, transport of nutrients, protein synthesis, neuroprotection, and apoptosis (Wesseling et al., 2020). However, aberrantly increased levels of soluble Tau protein often leads to intracellular toxicity. Likewise, excessive phosphorylation results in the self-assembly of tangles and straight filaments, which are involved in the physio-pathogenesis of AD-related neurodegenerative tauopathies, such as frontotemporal dementia (FTD-tau), progressive supranuclear palsy (PSP), and corticobasal degeneration (CBD) (Lathuilière et al., 2017).

Recent proteomics studies have identified over 50 candidate Tau phosphorylation sites during Tau aggregation into neurofibrillary tangles, supporting a critical role of Tau PTMs in AD progression (Barthélemy et al., 2019; Wesseling et al., 2020). The recent discoveries of Tau hyperphosphorylation sites have led to the development of biomarkers for the diagnosis of Tau-associated diseases. Indeed, Tau hyperphosphorylation at residues T153, T205, S208 and T217 were exclusively found in CSF from AD patients (Barthélemy et al., 2019). Also, increased plasma levels of p-Tau (T181) were used to stratify patients with AD dementia from non-AD neurodegenerative diseases (Janelidze et al., 2020). The hyperphosphorylation of residues T181 and T217 was shown to occur decades before the formation of NFT and onset of clinical symptoms (Barthélemy et al., 2020). And t-Tau and p-Tau (T205) were associated with brain atrophy, cognitive decline, and late onset of symptoms. In addition, hyperphosphorylation at Y18 residue is increased in the cortex at Braak stage V/VI in AD patients (Neddens et al., 2018). Those clinical observations indicate that phosphorylation alterations in the Tau protein follow a strict curve over the course of AD progression (Barthélemy et al., 2020). Genetically, several mutations in the Tau gene have been shown to cause neurodegenerative disorders including familial frontotemporal dementia (FTD) and progressive supranuclear palsy (PSP) and parkinsonism linked to chromosome 17 (FTDP-17) (Lopez Gonzalez et al., 2016; Morishima-Kawashima et al., 2011). The well-studied mutated Tau^P301L^ was first diagnosed in six French families with frontotemporal dementia and parkinsonism (Dumanchin et al., 1998), and has since been found in approximately 32 families around the world (Tsuboi et al., 2004; Kowalska et al, 2009). This proline to leucine substitution causes Tau aggregation and develops numerous intracytoplasmic tau deposits in multiple brain regions, including the hippocampus, neocortex, and substantia nigra (Giménez-Llort et al., 2008).

Vascular dementia (VD) is the second most common subtype of dementia after AD, accounting for 15-30% of dementia cases globally (Lobo et al., 2000; Rizzi et al., 2014; Jhoo et al., 2008; Chan et al., 2013; Kalaria et al., 2008). Recent epidemiological studies have established a link between AD and cardiovascular disease risk factors (Lamar et al., 2020) (e.g., hypertension, atherosclerosis, obesity, type 2 diabetes, etc.) suggesting that the neurovascular units (NVU) may contribute to the initiation and progression of AD (Zlokovic et al., 2011; Kim et al., 2020). These emerging data suggest that AD is a multifactorial, multi-organ disease with triggers beyond the CNS unit including vascular dysfunction and inflammation, which may precede neuropathological changes (De la Torre et al., 2018). In this context, Vascular smooth muscle cells (VSMCs) may play fundamental roles in regulating disease processes underlying both cardiovascular disease (CVDs) and AD (Frismantiene et al., 2018 and Mok et al., 2006).

VSMCs are highly plastic cells in the vessel wall (Castro et al., 2018; Rong et al., 2003) and readily transition from a contractile state to a synthetic or transdifferentiated entity in response to a wide range of pathogenic stimulus (Wang et al., 2015; Owens et al., 1995; Kalaria, 2003; Erkinjuntti et al., 2004). Transdifferentiated VSMCs are often characterized by loss of expression in contractile markers which are required to maintain the healthy vascular tone of brain arterioles. During vessel injury, VSMC phenotype switching is induced by various pro-inflammatory cytokines (i.e., interleukins and TNFα) (Tyrrell et al., 2021) or growth factors (i.e., PDGFs and FGFs) (Claesson-Welsh., 1996; Kendrick et al., 2011). Interestingly, synthetic VSMC-secreted proteins are associated with known neurogenerative markers, including the production of Coffilin-1, matrix metallopeptidase 9 (MMP9) and proinflammatory signals, such as CD68, IL-6, VCAM-1, ICAM-1 and MCP-1 (Lino Cardenas et al., 2019; Lee et al., 2013; Li et al., 2020). Therefore, there is a critical need to resolve the causal role of VSMCs and vascular dysfunction underlying AD progression. In this study we show that dysfunctional VSMCs can contribute to the AD development through the production of neuroinflammatory molecues including CD68, MMP9 and IL-6. We also show that VSMCs can generate elevated quantities of Tau protein as well induce its hyperphosphorylation at multiple sites, further implicating VSMCs in the initiation and progression of AD pathogenesis.

## RESULTS

### Dysfunctional VSMCs contribute to brain inflammation and Tau hyperphosphorylation

The role of brain-VSMCs in neuroinflammation and Tau hyperphosphorylation is poorly understood. Thus, we performed a transcriptomics analysis of previously described markers of VSMC phenotypic state from healthy and AD human brains (Bennett et al., 2014). Interestingly, AD-brains exhibited an up-regulation of synthetic-VSMC genes including *GJA, S100A4, CALD1* and *PDGFRB* (known to be a potent inducer of VSMC phenotype switching). Remarkably, genes that associated with vascular-related diseases including *LTBP3* (aortic dissection) (Guo et al., 2018), *PRKG1* (thoracic aortic aneurysm) (Isselbacher et al., 2016) and *TLN1* (coronary atherosclerosis) (Li et al., 2020) were also found to be up-regulated in AD-brains (Figure 1A and S1A). In contrast, expression of VSMC contractile genes in AD-brains were almost undetectable consistent with previous observations in other forms of VSMC-related diseases (Frismantiene et al., 2018; Owens et al., 2004; Mack, 2011). To investigate the role of VSMC phenotypic switching in neuroinflammation, we evaluated the levels of contractile proteins (SM22α) in arterioles and microvessels from brains of the humanized animal model of tauopathy and Alzheimer’s disease, the 3xTg-AD mice at 23 weeks (Oddo et al., 2003). Notably we found that leptomeningeal and cortical arterioles of 3xTg-AD brains had abnormal wall thickness with lower levels of Sm22α, which was associated with increased levels of platelet-derived growth factor-BB (PDGFBB), a high affinity ligand for PDGFRβ (Figure 1B). Furthermore, 3xTg-AD brains contained high levels of total Tau protein especially in brain arterioles as assessed by co-localized with Sm22α-positive VSMCs (Figure1C). To investigate whether transdifferentiation of VSMCs contributes to neuroinflammation, we treated brains from healthy mice with PDGFBB for 48hr (*ex vivo*) and analyzed markers of neuroinflammation by immunofluorescence. Our results showed that treated brains have elevated markers of neurotoxicity, including total Tau and CD68 (Figure 1D). Further, western blot analysis of total brain lysates with PDGFBB treatment showed a significant increase in p-Tau-S262 levels compared to control brains (DMSO, p<0.001) (Figure 1E). Quantitative proteomics analysis of PDGFBB versus DMSO treated brains revealed several AD-associated proteomic changes in these brains (Figure 1F). Pathway analyses of the proteomic data identified upregulation of multiple known AD pathways, including altered cell-to-cell signaling, neurological diseases, cell proliferation and neuroinflammatory pathways (Figure 1G). Based on these findings we then investigated the contributory portion of VSMCs in neuroinflammation and toxicity during AD pathogenesis. For this experiment, we genetically labeled brain VSMCs of wild-type mice with a red fluorescent protein (TdTomato) expressed under the control of the *Tagln* (Sm22α) promoter (Figure 1H). Then we subjected *Tagln-cre/*TdTomato brains to 3 *in vitro* models of neuroinflammation (PDGFBB, mutant Tau^P301L^ and LPS) to mimic AD-like microenvironments. Flow cytometry analysis of sorted VSMC^TdTomato^ from healthy brains revealed that vascular smooth muscle cells adopt a secretory phenotype compared to the DMSO controls (Figure 1I). We then quantified the inflammatory response of VSMC^TdTomato^ under AD-like conditions by flow cytometry using antibodies against CD68 (inflammation marker) and IL-6R (interleukin-6 receptor, a pro-inflammatory cytokine receptor). Our results demonstrate that the mutant Tau^P301L^ model had the highest proportion of CD68 and IL-R positive cells, followed by PDGFBB treatment. In contrast, LPS treated brains had the lowest proportion of both proinflammatory markers indicating a strong association of dysfunctional VSMCs in the inflammatory response rather than the activation of inflammatory pathways by LPS (Figure 1J). Taken together these findings provide new insights into the critical role for VSMCs in neurotoxicity.

**Figure 1.**
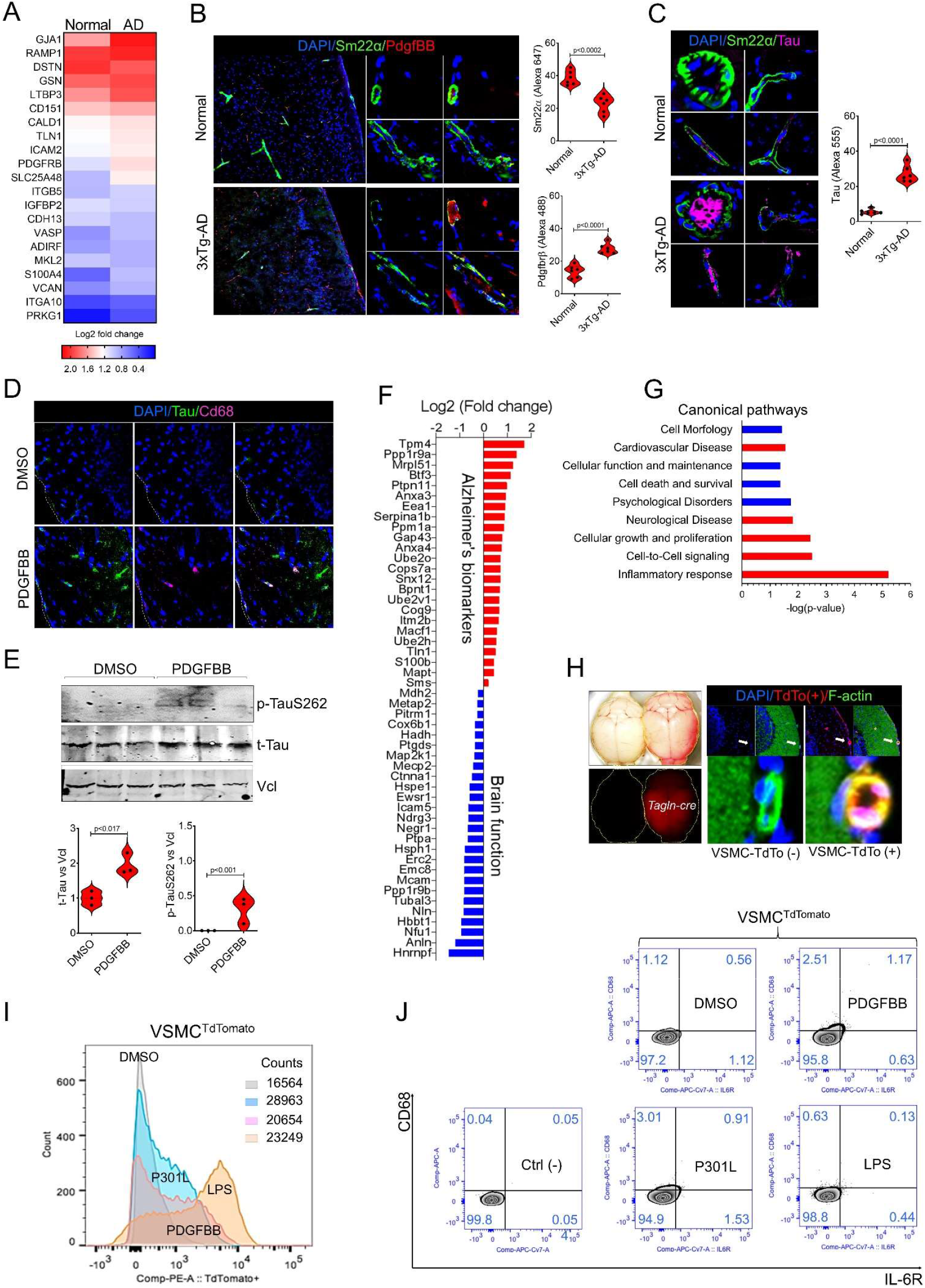
Dysfunctional VSMCs contribute to brain inflammation and Tau hyperphosphorylation. **(A)** Heatmap of differentially expressed VSMC marker genes (differentiated and modulated SMC) in post-mortem brains from non-demented control individuals (n=151) and AD patients (n=257). The scale bar indicates log2 fold change in expression determined from the limma R package. **(B)** Immunofluorescence staining of brain tissue from normal (n=6) and 3xTg-AD (n=6) mice at 22wks old, demonstrates loss of Sm22α (green) and increase in PdgfBB (red) in brain arterioles of the 3xTg-AD mice. **(C)** Immunofluorescence staining demonstrates accumulation of Tau protein (magenta) in brain arterioles (green) in brains of the 3xTg-AD mice. **(D)** Immunofluorescence analysis of Tau protein (green) and CD68 (magenta) in brains from healthy mice treated with DMSO (n=3) or PDGFBB (n=3). **(E)** Western blot demonstrates increased phosphorylation of the Tau-S262 and total Tau in brains treated with PDGFBB. **(F)** Proteomic signature of healthy brains treated with DMSO or PDGFBB. **(G)** Ingenuity path analysis (IPA) of biological pathways impacted by proteomic changes in brains treated with PDGFBB vs DMSO. **(H)** Generation of brain-VSMCs expressing fluorescent red TdTomato protein *in vivo*. **(I)** Representative histogram showing numbers of VSMC^TdTomanto^ from brains treated with PDGFBB (n=3), Tau^P301L^ (n=3), LPS (n=3) or DMSO (n=3). **(J)** Dot plot of healthy brains expressing VSMC^TdTomanto^ stimulated with PDGFBB, Tau^P301L^, LPS or DMSO by flow cytometry analysis.

### Morphological and phenotypic changes of VSMCs under AD-like conditions *in vitro*

To further investigate the impact of dysfunctional VSMCs in AD disease, we used primary human VSMCs from neural crest lineage (Steinbach et al., 2016) and primary human microglia cells. The human VSMCs that were treated with PDGFBB, mutant Tau^P301L^ and LPS showed changes in their cytoskeleton morphology characterized by the formation of dendritic-like phenotype (lamellipodium) and loss of polarized cytoskeleton microfilaments (F-actin) (Figure 2A, top panel). These changes were associated with increased Tau and MMP9 protein expression in VSMCs under AD-like conditions (Figure 2A, lower panel). Evaluation of VSMC phenotypic markers demonstrated a significant downregulation of contractile genes including *TAGLN* (SM22α), *LRP1, CNN1*, and consistent upregulation of matrix degradation and inflammatory markers such as *MMP2, MMP9, IL-6* and *CD68* (Figure 2B). Like our *ex vivo* results, flow cytometry analyses of VSMCs under AD-like conditions show elevated inflammatory markers (CD68 and IL-6R) in PDGFBB and Tau^P301L^ but not with LPS treatment compared with DMSO controls (Figure 2C), suggesting that VSMCs undergo cellular reprograming, and trans-differentiation into pro-inflammatory cells to promote neuroinflammation. As chronic inflammation has been shown to trigger microglial activation (Navarro et al., 2018; Hansen et al., 2018), we also investigated the response of microglial cells to transdifferentiated VSMCs using a wound healing assay. Our results showed that under normal conditions (DMSO) microglial cells have a propensity to inhibit the migratory properties of synthetic VSMCs. However, under AD-like conditions (PDGFBB, mutant Tau^P301L^ or LPS) microglia cells lose this ability while synthetic VSMCs displayed higher proliferation and migration activities (Figure 2C). Together these data indicates that AD brains favor the reprograming of VSMCs and may alter the physiological responses of vascular tissue thereby accelerating neurotoxicity in AD brains.

**Figure 2.**
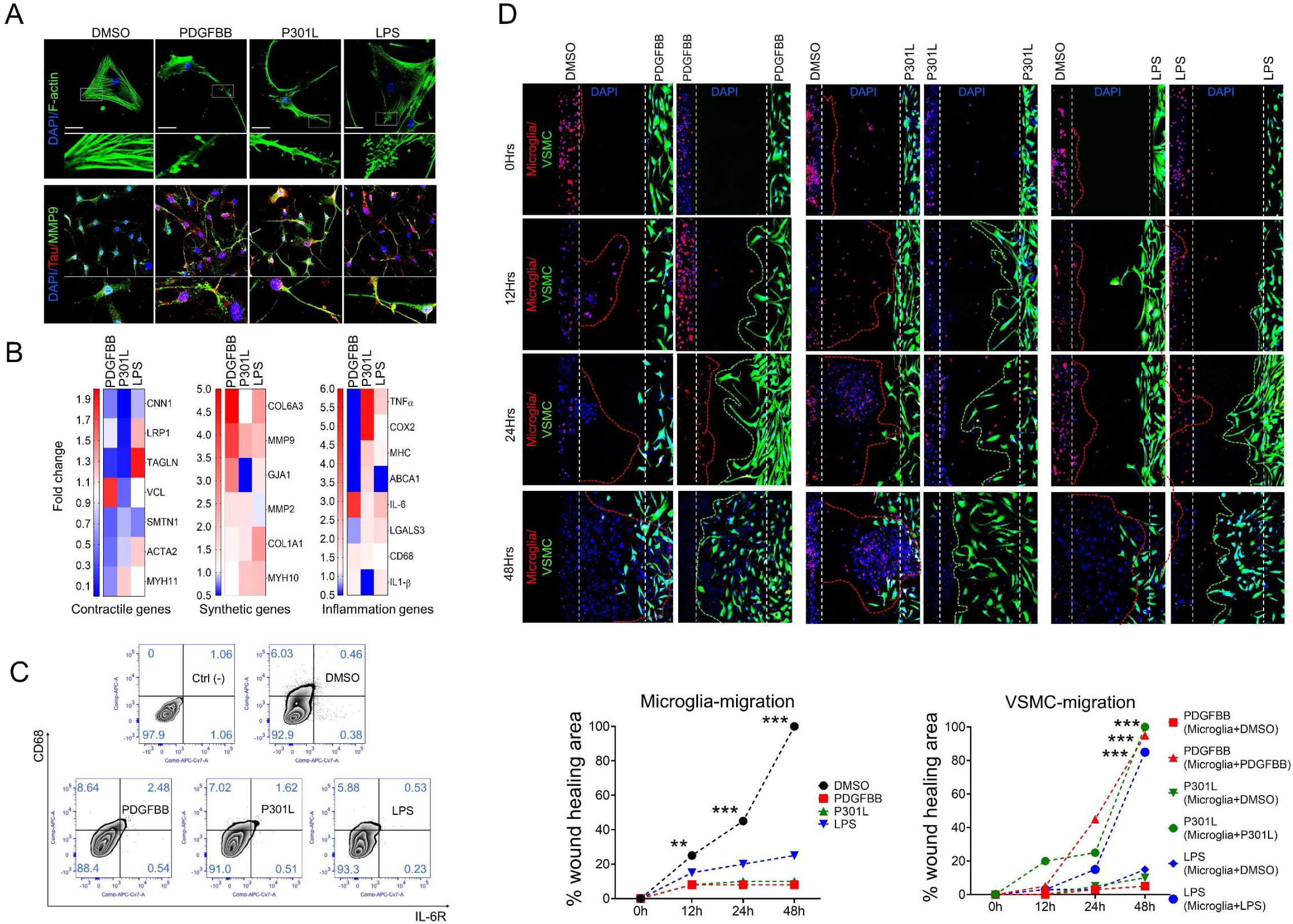
Morphological and phenotypic changes of VSMCs under AD-like conditions *in vitro*. **(A)** Immunofluorescence staining of primary human VSMCs from carotid arteries stimulated with PDGFBB, Tau^P301L^, LPS or DMSO, demonstrate morphologic changes in the cytoskeleton (F-actin, green, top panel) and increased Tau (red) and MMP9 (green) proteins (lower panel). **(B)** Heatmap of mRNA expression of contractile, synthetic, and inflammatory genes from VSMCs under AD-like conditions versus control group (DMSO). Red indicate upregulated, white indicate no significant change and blue indicate downregulated. **(C)** Flow cytometry analysis of CD68 and IL-6R expression in VSMCs under AD-like conditions. **(D)** Wound healing assay demonstrate that AD-like conditions favor the proliferative phenotype of VSMCs.

### Dysfunctional VSMCs induce AD-associated microglial phenotype

Next, we investigated whether dysfunctional VSMCs can activate microglial cells under AD-like condition. Therefore, we used a trans-well assay to quantify the migratory potential of microglia in response to the inflammatory signals from synthetic VSMCs treated with PDGFBB, Tau^P301L^ and LPS. To achieve this goal, we first labelled microglia cells with a membrane binding red fluorophore under normal conditions and in parallel VSMCs were plated under AD-like conditions for 72hr in a 24 trans-well plate followed by incubation with growing media for 16hr. Remarkably, microglial cells migrated significantly towards to VSMCs that were exposed to PDGFBB, mutant Tau^P301L^ and LPS (Figure 3A, top panel). Immunofluorescence analysis of pro-inflammatory markers for activated microglia (M1 microglia) such as MHC-II and iNOS demonstrated that under AD-like conditions, VSMCs can induce the activation of microglia potentially through a paracrine communication. Interestingly, confocal analyses revealed CD68 colocalization with SM22α, with evidence of microglia physically interacting with VSMCs under AD-like conditions, analogous to previously reported microglia-neuron interactions in the brain (Pósfai et al., 2019) (Figure 3A, middle panel). In contrast, pre-treatment of microglia cells with all three AD-like conditions failed to promote the migration of healthy VSMCs (Figure 3A, Lower panel). Immunostaining showed increased levels of MMP9 and PDGFR-β in VSMCs under AD-like conditions (Figure 3B). However, PDGFBB treatment failed to induce MMP9 expression in microglial cells, supporting the notion of a paracrine VSMC-mediated activation of microglial cells in AD brains (Figure 3B). To support this idea, we performed a co-culture experiment by pre-treating healthy VSMCs with PDGFBB, mutant Tau^P301L^ and LPS for 72hr followed by co-culture with healthy microglia cells in normal conditions. Indeed, VSMCs pre-treated with PDGFBB and Tau^P301L^ induced expression of the classic M1 microglial marker, MHC-II but, LPS-treated VSMCs failed to activate microglia (Figure 3C). Then we performed a second experiment using the supernatant of VSMCs pre-treated with all 3 AD-like conditions to incubate healthy microglia cells for 48hrs. Immunofluorescence analysis showed that supernatants from VSMCs treated with PDGFBB and Tau^P301L^ was sufficient to promoted morphologic and phenotypic changes corresponding to activated microglia associated with brain neuroinflammation in AD patients (Figure 3D) (Leng at al., 2020)al. Collectively, our results suggest that dysfunctional VSMCs can activate microglia to drive brain neuroinflammation.

**Figure 3.**
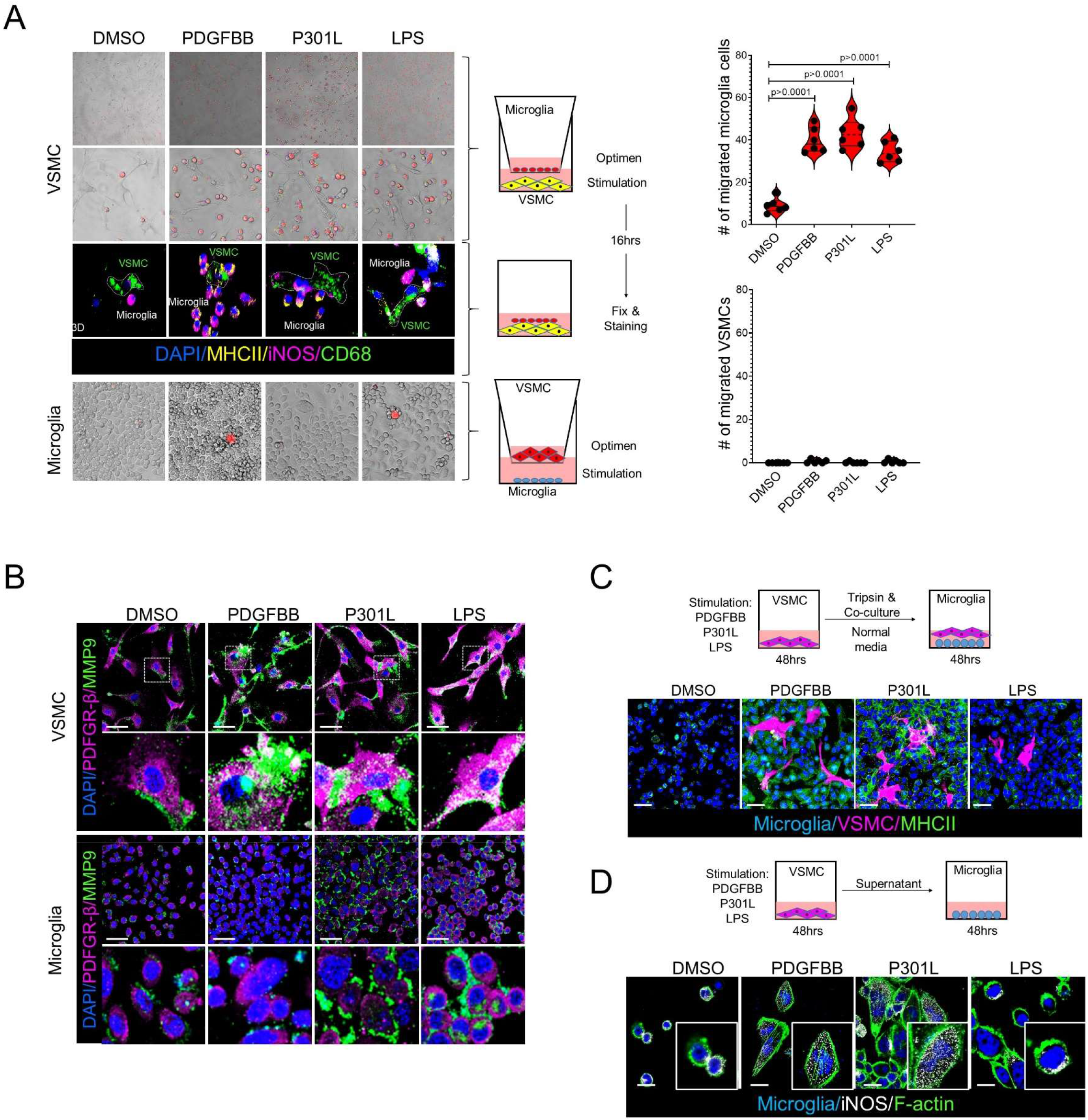
Dysfunctional VSMCs induce AD-associated microglial phenotype. **(A)** Representative trans-well migration and invasion assay shows enhanced trans-well migration of microglia towards to VSMCs under AD-like conditions (top panel). In contrast, microglia under AD-like conditions did not promote trans-well migration of healthy VSMCs. **(B)** Immunofluorescence staining of PDGFBR-β (magenta) and MMP9 (green) in VSMCs and microglia under normal (DMSO) or AD-like conditions. **(C)** Representative micrograph of co-culture assay of VSMCs (magenta) stimulated with PDGFBB, Tau^P301L^, LPS or DMSO and healthy microglia cells (blue). **(D)** Immunofluorescence staining of microglia cells incubated with supernatants from VSMCs stimulated with PDGFBB, Tau^P301L^, LPS or DMSO, shows induction of microglia activation assessed by iNOS staining.

### *In silico* analysis of mutant Tau proteins and *in vitro* and *in vivo* Tau hyperphosphorylation status

To better understand the molecular and structural impact of mutations in the Tau protein associated with neurodegenerative diseases, we performed an *in-silico* study using molecular dynamics simulations (MDS) (Table S1). By using the Tau full-length 2N4R sequences (441 aa), we built the 3D structure of the protein (Figures S2A and S2B. Movie S1). Then based on this 3D model, the Tau^P301S^ and Tau^P301L^ were built performing 301 site direct mutagenesis using UCSF Chimera software (Pettersen et al., 2004), allowing local structural relaxion to avoid steric residue clashes (Figure 4A and S2C). Tau proteins are considered intrinsically disordered, there is evidence that in solution, they can adopt multiple globular structures (Popov et al., 2019). Thus, we sought to analyze all three Tau proteins with 500 ns of MDS to stabilize the physical motions of atoms in the Tau proteins and thus mimic physiological conditions. Importantly the residue 301 is located at the microtubule binding repeats domains R2 and R3 (MTBRs) where the assembly and stabilization of axonal connections occurs at the neuronal cell body (Figure S3B). Even though all three proteins fold towards a globular structure, the folding was remarkably different for the Tau protein with a proline to leucine substitution compared to the proline to serine substitution, in terms of folding patterns. While looking at the global surface of the Tau^P301L^, we observed a decrease in the solvent-accessible surface area (SASA) from 360 nm^2^ to 240.31 nm^2^. The MTBR domain showed a significant increase in the SASA (7%), potentially exposing a greater area of interaction thus creating a predisposition to posttranslational modification including phosphorylation of multiple residues (Table S2). Another important consequence of the proline to leucine substitution was observed on the electrostatic potential (ESP) surfaces at the MTBRs. All three proteins show nucleophilic and hydrophobic interactions. However, the R2-R3 domains of the Tau^P301L^ adopt an additional negative electrostatic pattern creating a more promiscuous Tau protein which may be more prone to interact with multiple Tau-protein kinases, ligands, and drugs (Figure S4A-S4C). To achieve these conformational changes the proline to leucine substitution induces loss of residue contacts at H299, V300, G303, G304, S305, V309, L325 and creates uncommon interactions with residues at Q307, K321 and N-terminal regions (Figures 4A, 4B, S3A and S3C-S3E). Remarkably, the misfolding of the Tau^P301L^ protein reduced the distance of the Y18 residue toward to the MTBRs right where the P301L substitution occurs (Figure 4C and S5E). In this context the S262 residue showed the largest area of exposure to the solvent and thereby augmenting its capability to interact with Tau-protein kinases (Figure S6A). Similarly, multiple Tau residues previously associated with brain neurodegeneration were relocated at the proximity of the proline to leucine substitution and augmenting their ability to posttranslational modifications (Figure 4B and Movie S2). In agreement with our *in silico* results, we observed that VSMCs under AD-like conditions produced high levels of total and phosphorylated Tau. Indeed, VSMCs treated with PDGFBB and Tau^P301L^ promoted predominately the hyperphosphorylation of the S262 residue and also other residues at Y18 and T205, which associated with up-regulation of Cofilin-1, a proliferative-VSMC marker (San et al., 2008). Interestingly, Cofilin-1 has recently been implicated in the pathogenesis of AD (Kang et al., 2019), supporting the potentially role of dysfunctional VSMCs in neurotoxicity and the AD progression (Figure 4D and S1A). Immunofluorescence of 3xTg-AD brains at 13 weeks and 22 weeks demonstrated that loss of contractile proteins (Sm22α and α-Sma) in brain arterioles occurs in an age-dependent manner which associated with an increased production of CD68 by VSMCs during the disease state (Figure 4E) Therefore, Tau misfolding is associated with hyperphosphorylation and strongly supports the notion that dysfunctional VSMCs can induce conformational changes in Tau protein and neurotoxicity previously implicated in tauopathies including AD.

**Figure 4.**
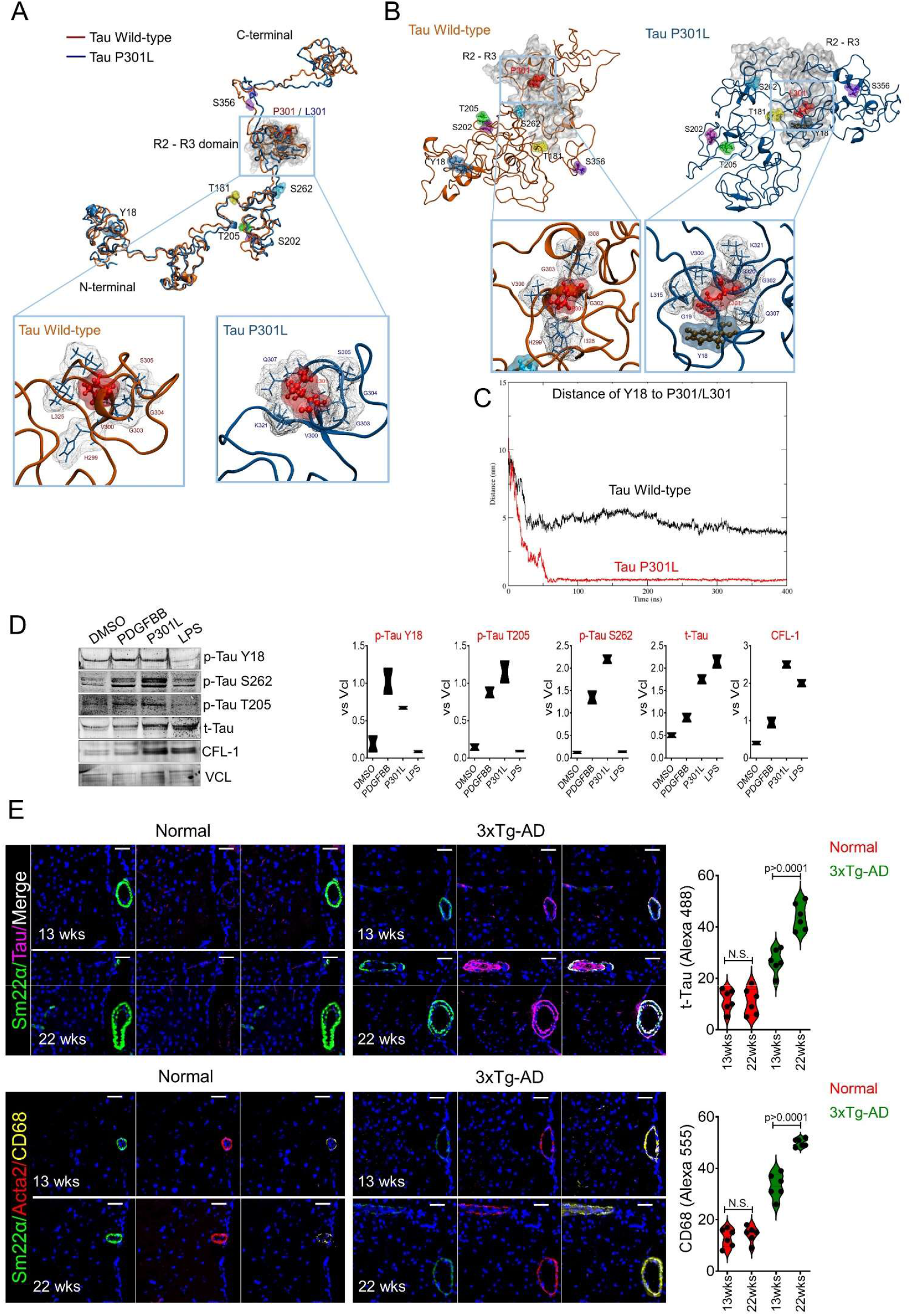
*In silico* analysis of mutant Tau proteins and *in vitro* and *in vivo* Tau hyperphosphorylation status. **(A)** Representative western blot image of levels of t-Tau and multiple phosphorylate Tau residues from total lysates of VSMCs under AD-like conditions **(B-C)** in silico analysis of the WT and mutant Tau^P301L^. **(D)** Immunofluorescence analysis of AD markers, Tau (Magenta), CD68 (yellow) and VSMC marker Sm22α (green) and Acta2 (red) in brain from normal and 3xTg-AD mice at 13wks and 22wks.

## DISCUSSION

Vascular dysfunction is a well-established clinical aspect of AD, with subtle changes in microvasculature of the brain manifesting at the onset of clinical symptoms (Govindpani et al., 2019, Arvanitakis et al 2016). However, the precise role of VSMCs in the pathogenesis of AD remains unclear. The neurovascular unit (NVU) is primarily composed of endothelial cells, pericytes, astrocytes and vascular smooth muscle cells (VSMCs) (Zlokovic et al., 2011). Given the inherent phenotypic plasticity of vascular smooth muscle cells, they readily switch from normal contractile to synthetic phenotypes in response to a variety of environmental stimuli, including inflammatory cytokines, genetic mutations or chemical compounds. Previous neuropathological studies of AD autopsies showed that 80% of AD patients without vascular dementia developed abnormal vascular features, including multiple micro-infarcts, lacunes, cerebral micro-bleeds indicative of intracranial atherosclerosis, perivascular spacing and cerebral amyloid angiopathy (CAA) (Toledo et al., 2013). Together these observations support the concept that cerebrovascular dysfunction is a prominent pathological feature of AD, and that the neurovascular unit is a central catalyst of AD initiation and progression.

Our findings point towards the contribution of dysfunctional VSMCs in the development of neuroinflammation and AD progression. In this study, downregulation of contractile genes critical for the homeostasis of brain arterioles was tightly linked to the production of neuroinflammatory markers such as MMP9, CD68 and IL-6. Moreover, transdifferentiated VSMCs can produce high levels of total and hyperphosphorylated Tau protein. Remarkably under AD-like conditions VSMCs promoted hyperphosphorylation of Tau-T205 which has been associated with early neurological decline in AD patients, further illuminating the contribution of VSMCs in the initiation of clinical symptoms. We found that brain arterioles of 3xTg-AD mice contained high levels of PDGFBB (Platelet-derived growth factor-BB) which is a well-known inducer of VSMC transdifferentiation through a potent mitogenic activity by inhibiting expression of contractile genes such as αSMA (*ACTA2*), SM22α (*TAGLN*), LRP1 and CNN1 (Calponin) (Tallquist et al., 2004; Salabei et al., 2013). Furthermore, MMP proteins known to be associated with other vascular smooth muscle cell related diseases including aortic aneurysm, vascular calcification, and vascular stenosis, was also found elevated in postmortem brain tissue of AD patients. In agreement with our work, elevated MMP proteins in VSMCs under AD-like conditions coincided with downregulation of SM22α, which is known to be a negative regulator of MMP9 (Nair et al., 2006) indicating the negative feedback regulation of neuroinflammatory molecules due to loss of VSMC-contractile phenotype. Similarly, downregulation of LRP1 in VSMCs was observed in AD-like conditions and may impact Aβ clearance leading to Aβ accumulation and plaque formation during diseases (DeSimone et al., 2017; Kanekiyo et al., 2012; Nelson et al., 2017). Along these lines, microglia cells have an increased phagocytic response to Aβ and are known effectors of Aβ clearance in the brain (Lopes et al., 2008; Streit et al., 2009). However, during AD, microglia adopt reactive states characterized by both morphological and functional changes including increased expression of iNOS and MHC-II and other acute inflammatory molecules (Zabel et al., 2013). Our *in vitro* assays demonstrated the potential of dysfunctional VSMCs to trigger AD-associated microglial phenotypes. These changes were more remarkable in PDGFBB and Tau^P301L^ conditions, further supporting the contribution of unhealthy VSMC in promoting AD related pathology features and neurodegeneration.

Some forms of genetically triggered neurodegenerative diseases such as Tau^P301L^ protein promote neurofibrillary deposits (Kawasaki et al., 2020). Tau hyperphosphorylation may promote self-aggregation and subsequent fibrillization (Eschmann et al., 2017; Berendsen et al., 2000). Our *in silico* analysis of the 3D structures of wild-type and Tau^P301L^ proteins demonstrates that changes in the amino acid sequence could have a dramatic impact on the conformational structure and misfolding, leading to the phosphorylation of multiple residues via rearrangement of key amino acids (e.g. Y18, T181, S202, T205, S262 and S356) involved in AD progression. In agreement with our *in silico* analysis, we found that dysfunctional VSMCs can induce Tau misfolding and promote its hyperphosphorylation. In addition, our *in vivo* studies reveal that brain arterioles contained significant levels of Tau protein and inflammatory markers co-localizing with VSMCs in 3xTg-AD mice. Additionally, we uncovered that Tau protein levels and inflammatory signal were relative to the age of the 3xTg-AD mouse brain indicating that the vascular tissue may play a pathological role in the development of the disease as early as before the outcome of clinical symptoms.

Overall, this study demonstrates the pivotal role of dysfunctional VSMCs in AD development and progression. It has refined our understanding of the contributory role of the neuro-vascular unit and the direct contribution of VSMC towards Alzheimer disease pathogenesis. While the majority of AD studies have focused on neural and glial cells, this study extends the catalog of promoting factors in AD which promise to enable development of new therapeutic interventions and vascular-related biomarkers for early detection of the disease. In this context, recent studies have showed that targeting the vascular tissue with drugs such as Losartan, ACEi or Captopril could help to improve memory and cognitive decline (Lee et al., 2020; Royea et al., 2020; Royea et al., 2020; Asraf et al., 2018; Rahimi et al., 2020). Therefore, this study supports the idea of VSMCs as a novel therapeutic target in AD.

## METHODS AND DETAILS

### Human Alzheimer Disease vascular SMC gene expression analysis

Bulk RNA-seq based gene expression data were analyzed from the dorsolateral prefrontal cortex (post-mortem) from n=414 individuals (n=151 cognitively normal, non-demented controls and n=263 AD subjects) in the Religious Orders Study and Rush Memory and Aging Project (ROSMAP) (PMID: 31340147). These data were downloaded from the AMP-AD Knowledge portal on the Synapse platform (Sage Bionetworks) using accession number: syn3388564 (https://doi.org/10.7303/syn3388564). Briefly, differentially expressed genes (DEG) were detected between non-demented controls and AD brains using the *limma* package, after normalizing the expression values and removing batch effects and adjusting for covariates: age at death, sex, postmortem interval and APOE risk allele status. Genes were ranked by log2 fold change and posterior error probability. DEGs were then intersected with differentiated and modulated vascular SMC marker genes previously identified from single-cell RNA-seq of human coronary artery tissues (n=34 subjects) (PMID: 31359001). These DEGs were also queried in single-nuclei RNA-seq of post-mortem human control and AD brain tissues (n=24 subjects), courtesy of Dr. Philip De Jager and Dr. Vilas Menon (unpublished data).

### Animals

All mice were cared under strict compliance with the Partners Institutional Animal Care and Use Committee (IACUC), regulated by the United States Department of Agriculture and the United States Public Health Service. Tagln-cre mice (B6.Cg-Tg(Tagln-cre)1Her/J, # 017491) and TdTomato mice (B6.129S4-Gt(ROSA)26Sortm1Sor/J, Stock No: 003474 | R26R). were acquired from Jackson laboratories. Fixed brains from 3xTg-AD mice (B6;129-Tg (APPSwe,tauP301L)1Lfa Psen1tm1Mpm/Mmjax, MMRRC Stock No: 34830-JAX), were acquired from Jackson laboratories.

### Cell lines

Primary human vascular smooth muscle cells from carotids of healthy donors were purchased from Cell Applications Inc. (#3514-05a). Human microglia were purchased from ScienCell Inc. (#1900). VSMC identity was assessed by immunofluorescence staining of contractile markers, including SM22α and CNN1. To preserve cell identity, all experiments were carried out at passages 1-5.

### AD-like models

For Ex-vivo experiments, brains from healthy mice were collected and immediately incubated in normal media (Cell Applications, #311-500) for 2 hrs at 37C and 5% CO_2_. Then, brains were treated with PDGFBB (n=3, 20ng/mL, Sigma-GF149), recombinant Tau P301L (n=3, 20µg/mL, rPeptide-T1014-1) or LPS (n=3, 0.8µg/mL, Sigma-L9143) for 48hrs, separately. For in vitro experiments, either VSMCs or microglia cells were treated with PDGFBB (n=4, 15ng/mL, Sigma-GF149), recombinant Tau P301L (n=4, 10µg/mL, rPeptide-T1014-1) or LPS (n=4, 0.5µg/mL, Sigma-L9143) for 72hrs, separately. Control brains (n=3) were treated with DMSO at 0.01%. Cells were supplied with growing media containing all three compounds every 24hrs, separately.

### Proteomic analysis

Approximately 400µg of total proteins were extracted from brains treated with DMSO (n=3), PDGFBB (n=3), recombinant Tau P301L (n=3) or LPS (n=3) using RIPA buffer (ThermoFisher, CA, USA) and supplemented with 1× of protease inhibitor cocktail (Roche) according to the manufacturer’s instruction, on ice for 30min. The protein concentration was determined by a BCA protein assay kit (Pierce Biothech), and then denatured protein samples were separated by SDS-PAGE followed by Coomassie staining. Then gel bands containing separated proteins were analyzed using an Orbitrap mass spectrometer (Thermo Scientific, CA, USa), as reported previously (Ibrahim et al., 2016). Ingenuity Pathway Analysis (IPA, Ingenuity Systems) software was used to identify potential canonical pathways associated with proteomic changes in brains under AD-like conditions.The predicted activation state and activation Z-score are based on the direction of fold change values for those genes in the input data set for which an experimentally observed causal relationship has been established (Kramer et al., 2013).

### Quantitative flow cytometry

For the ex-vivo brains were homogenized and stained with anti-mouse CD68 (clone FA-110) and anti-mouse IL-6R (clone D7715A7) BioLegend. APC-conjugated anti-Feeder Cells (clone mEF-SK4) was purchased from Miltenyi Biotec. Cells were surface stained by incubation with the relevant antibodies diluted in PBS + 2% FBS for 20 minutes at room temperature, followed by 2 washes with PBS 2% FBS. When intracellular staining of signature cytokines was performed, cell suspensions were initially incubated for 3–5 hours at 37°C with RPMI T cell media containing 0.1% ionomycin (MilliporeSigma, I0634), 0.1% brefeldin A (BioLegend, 420601), 0.1% monensin (BioLegend, 420701), and 50 ng/ml of PMA (MilliporeSigma, P8139). After the incubation, surface staining was performed as indicated above, followed by cell fixation for 20 minutes at room temperature in light-protected storage with Fixation buffer (BD Biosciences, 554655). Upon fixation and being washed with PBS 2% FBS, cell suspension was permeabilized with 1× wash/permeabilization buffer (BD Biosciences, 554723; diluted in distilled water) and intracellular stained for 20 minutes at room temperature in light-protected storage. Absolute cell numbers were quantified using Precision Count Beads (BioLegend). Flow cytometry data was acquired on a LSRII (39) and analyzed using FlowJo software.

### QPCR

Total RNA was extracted from VSMCs under AD-like conditions using miRNeasy kit (Qiagen) following the manufacturer’s protocol. RNA samples were used to hybridize Agilent gene expression microarrays (v2. mRNA microarrays, Agilent). Briefly, 100 ng of total RNA was used as the starting template for cDNA synthesis. This cDNA was used as a template to synthesize Cy3-labeled cRNA that was hybridized on 8 × 60 K high density SurePrint G3 gene expression human Agilent microarrays at 65 °C for 17 h. All arrays were normalized by cyclic-LOESS and significance analysis of microarrays (SAM) was applied for statistical analysis of the differentially expressed genes setting a 5% false discovery rate as a cutoff of statistical significance. 44 genes were found to be dysregulated in the same direction for both siRNA conditions compared with the control group. Fourteen genes were randomly chosen to be validated by qRT–PCR. RNA samples were used to run qRT–PCR on SYBR green system (Applied Biosystem, Foster city, CA). Results were analyzed by the ddCT method and GAPDH (encoding glyceraldehyde-3-phosphate dehydrogenase) was used as a housekeeping gene. Fold change were calculated by taking the average over all the control samples as the baseline.

### Wound healing assay

Human VSMCs and microglial were seeded into silicone inserts plates with a defined cell-free gap (Ibidi-labware) and transfected 24hrs with specific stimulation. After 48 h silicone inserts were removed to generate the initial wound. No antiproliferative agents were added to the media, therefore wound healing in our assay reflects the combined activity of migration and proliferation. The wound healing/closure was evaluated 12hrs, 24hrs and 48hrs later.

### Cell migration and invasion assay

Briefly, 500,000 VSMCs were seeded on the bottom and treated with PDGFBB (15ng/mL, Sigma-GF149), recombinant Tau P301L (10µg/mL, rPeptide-T1014-1) or LPS (0.5µg/mL, Sigma-L9143) for 72hrs, separately. And 300,000 microglia were labeled with a cell-permeable dye (ThermoFisher, #R37601) and seeded on the membrane insert under normal conditions. Then both cell lines (VSMCs-Bottom and microglia-Top) were incubated in normal media for 16 hrs. Next the membrane insert was removed and cells in the bottom of the plate were fixed with 4% formaldehyde for quantification and immunofluorescent staining. 24 transwell plates (pore size, 8µm) were used according to the manufacturer’s instructions (Corning, CLS3464-48EA).

### Immunoblotting

Nuclear and cytoplasmic protein lysates were prepared using NE-PER Kit (Pierce, Rockford, IL, USA) and supplemented with 1× of protease inhibitor cocktail (Roche) according to the manufacturer’s instruction. For detection of nuclear proteins 20 µg of total nuclear extracts were mixed with denaturing buffer (1× Laemmli loading buffer with 10% of β-mercaptoethanol) and analyzed by SDS–PAGE/western blot. Separated proteins were transferred onto a nitrocellulose membrane using the iBlot transfer system (Novex, ThermoFisher, USA). For detection of cytoplasmic proteins 15 µg of cytoplasmic extracts were mixed with denaturing buffer as described above. In general, primary antibodies were used at concentration of 1:100 and secondary at concentration of 1:10000, although specific concentrations are listed in Supplementary Data 7.

### Computational Details

#### System building, structural refinements and molecular dynamic simulations (MDS)

The 2N4R Tau sequence (441 aa), was used to build the wild-type Tau protein 3D structure using the I-TASSER server (Roy et al., 2010; Yang et al., 2015; Yang and Zhang, 2015). The TauP301S and TauP301L were built based on wild-type 3D model by performing 301 site direct mutagenesis using UCSF Chimera software (Pettersen et al., 2004). Then the structural refinement was carried out to avoid residues overlapping using ModRefiner server (Xu and Zhang, 2011). Classical MD simulations were performed using GROMACS 2020.4 package (Abraham et al., 2015) with the OPLS-AA force field parameters (Jorgensen et al., 1996). All three Tau protein systems were built in a triclinic simulation box considering periodic boundary conditions (PBC) in all directions (x, y, and z) (Table S1). Then, they were solvated using the TIP4P water model (Jorgensen et al., 1985), and Cl^-^ ions were used for neutralization of total charge in the simulation box. Mimicking of physiological conditions was performed by ionic strengthening with the addition of 150mM NaCl. The distance of the protein surfaces to the edge of the periodic box was set at 1.5nm. And 1 fs step was applied to calculate the motion equations using the Leap-Frog integrator (Hockney et al., 1974). The temperature for proteins and water-ions in all simulations was set at 309.65 K using the modified Berendsen thermostat (V-rescale algorithm) (Berendsen et al., 1984) with a coupling constant of 0.1 ps. Pressure was maintained at 1bar using the Parrinello-Rahman barostat (Bussi et al., 2007) with a compressibility of 4.5×10^−5^ bar^-1^ and a coupling constant of 2.0 ps. Particle mesh Ewald method (Ewald, 1921) was applied to long-range electrostatic interactions with a cutoff equal to 1.1nm for non-bonded interactions with a tolerance of 1×10^5^ for contribution in real space of the Tau 3D structures. The Verlet neighbor searching cutoff scheme was applied with a neighbor-list update frequency of 10 steps (20 fs). Bonds involving hydrogen atoms were constrained using the Linear constraint solver (LINCS) algorithm (Hess, 2008). Energy minimization in all simulations was performed with the steepest descent algorithm for a maximum of 100,000 steps. For the equilibration process, we performed two steps, one step of dynamics (1ns) in the NVT (isothermal-isochoric) ensemble followed by 2 ns of dynamics in the NPT (isothermal-isobaric) ensemble. Then the final simulation was performed in the NPT ensemble for 500ns followed by the analysis of the Tau 3D structures and their energy properties.

### Structural and energetic analysis of Tau 3D protein structures

All MD trajectories were corrected, and the Tau 3D structures were recentering in the simulation boxes. RMSD, RMSF, radius of gyrations, H bonds, residue distances, and solvent accessible surface area analyses were performed using the Gromacs tools and results were plotted using XMGrace software (Turner, 2005). For visualization of Tau 3D structures, we used the UCSF Chimera, VMD software packages (Humphrey et al., 1996). Analysis of atomic interactions and 2D plots were generated using the LigPlot software packages (Wallace et al., 1995). The electrostatic potential (ESP) surfaces were calculated using the APBS (Adaptive Poisson-Boltzmann Solver) software, and the PDB2PQR software was used to assign the charges and radii to protein atoms (Baker et al., 2001; Dolinski et al., 2004). FEL maps were generated to visualize the energy associated with the conformational structure of all three Tau proteins during the molecular dynamic simulation. FEL maps were generated using the RMSD and radius of gyration as atomic position variables respect to its average structure. FEL maps were plotted using using gmx sham and figures were built using Wolfram Mathematica 12.1 (Wolfram Research, Inc., 2020).

### Calculation of binding free energy

The Molecular Mechanics Poisson-Boltzmann Surface Area (MM/PBSA) calculation of free energies and energy contribution by individual residues were carried out to analyze the impact of amino-acid substitutions (P301S and P301L) in the R2-R3 domains using the last 50ns of MD trajectories and the g_mmpbsa package (Kumari et al., 2014). Therefore, the interacting energy was calculated using the following equation:

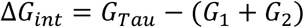

Where the terms G_1_ and G_2_ are the free energy of the different sites of the Tau protein, and G_Tau_ is the free energy of entire Tau 3D structure. In this context, the free energy of each term was calculated as follow:

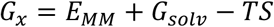

where E_MM_ is the standard mechanical energy (MM) produced from bonded interactions, electrostatic interactions, and van der Waals interactions. G_solv_, is the solvation energy that includes the free energy contributions of the polar and nonpolar terms. In this work, we used the solvent accessible volume (SAV) model to calculate the nonpolar term. The TS term refers the entropic contribution and was not included in this calculation due to the computational costs (Brown et al., 2009; Rastelli et al., 2010; Kumari et al., 2014). Finally, 309 Kelvin (K) of temperature was used as the default parameter in all our calculations.

### Statistical analysis

Results are given as mean ± SD. Student’s test was applied to determine the statistical significance of difference between control and treated groups (*p < 0.05, **p < 0.01 and ***p < 0.001). For all experiments at least three replicates were performed. Two-way analysis of variance (ANOVA) was used to analyze healing versus time of Figure 2D (95% confidence interval is plotted). P-values represent one-way ANOVA followed by Tukey’s honestly significant difference (HSD) post-hoc test. All graphs were produced using GraphPad Prism 8.0.

## SUPPLEMENTARY LEGENDS

**Figure S1. Tau protein structures used in this work**.

**(A)** Circos plot of the full-length Tau protein (2N4R) linear sequence.

**(B)** Front and top views of the wild-type Tau 3D protein obtained with I-Tasser server.

**(C)** Close up of the R2-R3 domain and alignment of Tau wild-type (orange color), Tau^P301L^ (blue), and Tau^P301S^ (green).

**Figure S2.**
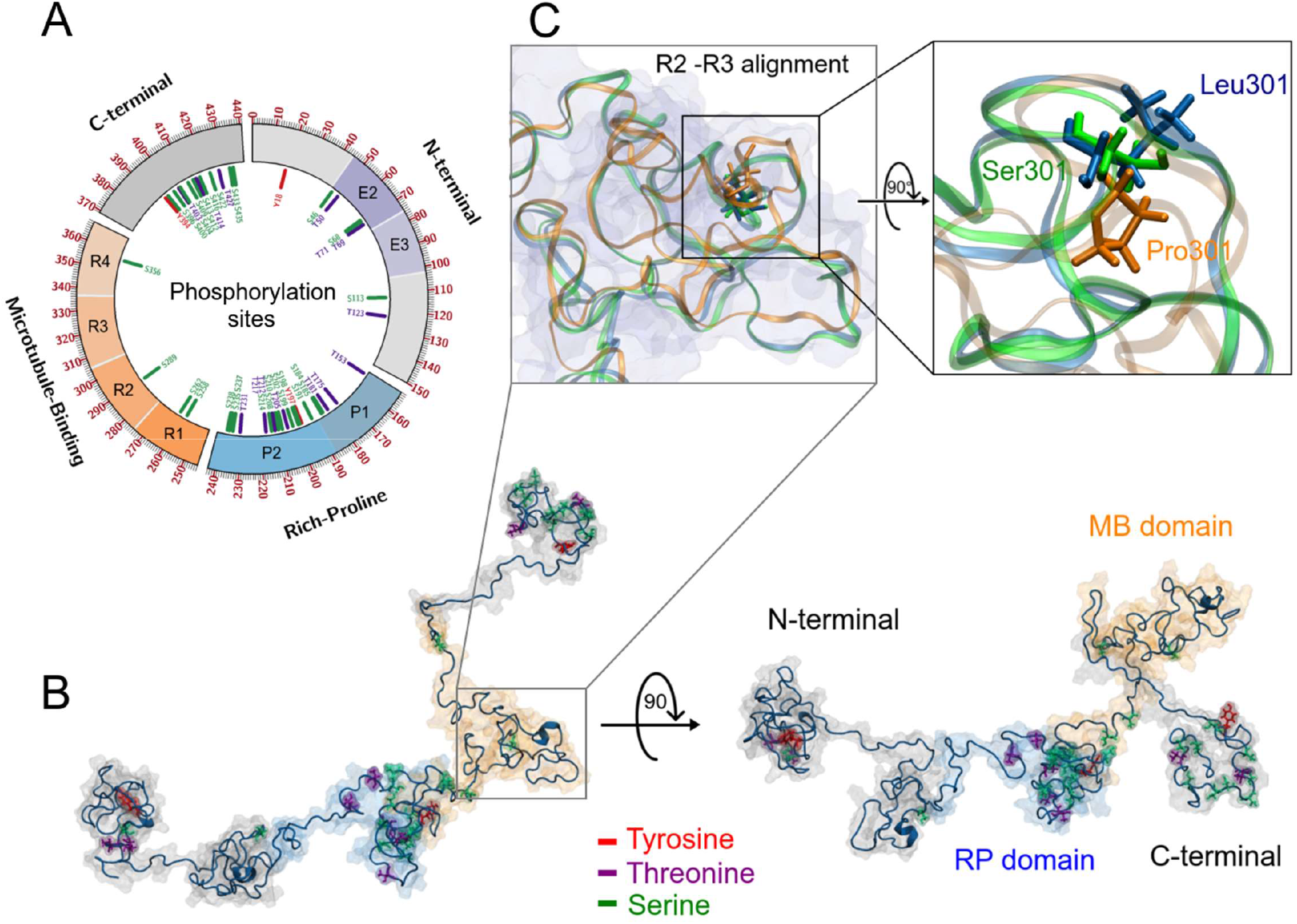
Effects of the 301 amino-acid substitution on the structural energy. **(A)** Plot of energy interaction of the residue 301 **(B)** Circular heat map of the vibrational frequencies and the β-factor. Red color indicates highest vibration and energy, green color indicates lowest fluctuation areas and yellow indicates moderate vibration and energy. The upper lines indicate vibrational frequencies of Tau wild-type (orange), Tau^P301S^ (green) and Tau^P301L^ (blue). The inner lines indicate interactions of the phosphorylated residues with their neighboring residues after 500ns of MD simulations. **(C, D, E)** MM/PBSA calculation of interacting environments for the Tau wild-type (orange), Tau^P301S^ (green) and Tau^P301L^ (blue). The 3D structures show the repulsion residues (red) and stable residues (blue). While the 2D maps show hydrophobic and electrostatic interaction at the 301 residues.

**Figure S3.**
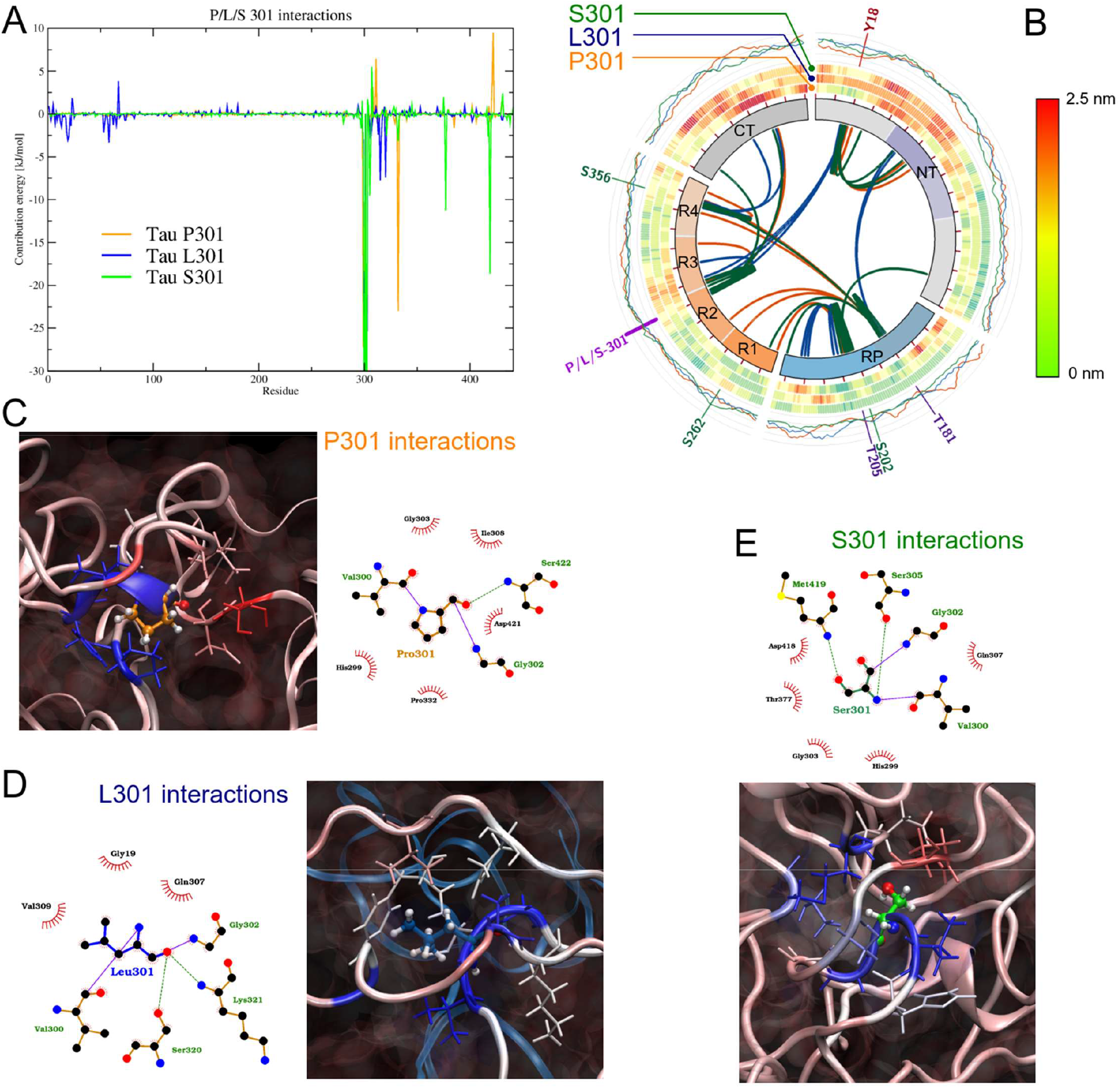
Electrostatic Potential (ESP) surfaces of the Tau proteins obtained at the final of molecular dynamic simulations. The red color indicates nucleophilic regions; the blue color indicates electrophilic regions; and the white indicates hydrophobic regions. The ESP units used was *k*_*B*_T/*e*, where *k*_*B*_ is the Boltzmann constant; T was the MDS temperature, and *e* was the electron charge. The ESP results were mapped on the protein surfaces at 2.0 isodensity values. **(A)** Tau wild type protein. **(B)** Tau^P301L^ protein, and **(C)** Tau^P301S^ protein.

**Figure S4.**
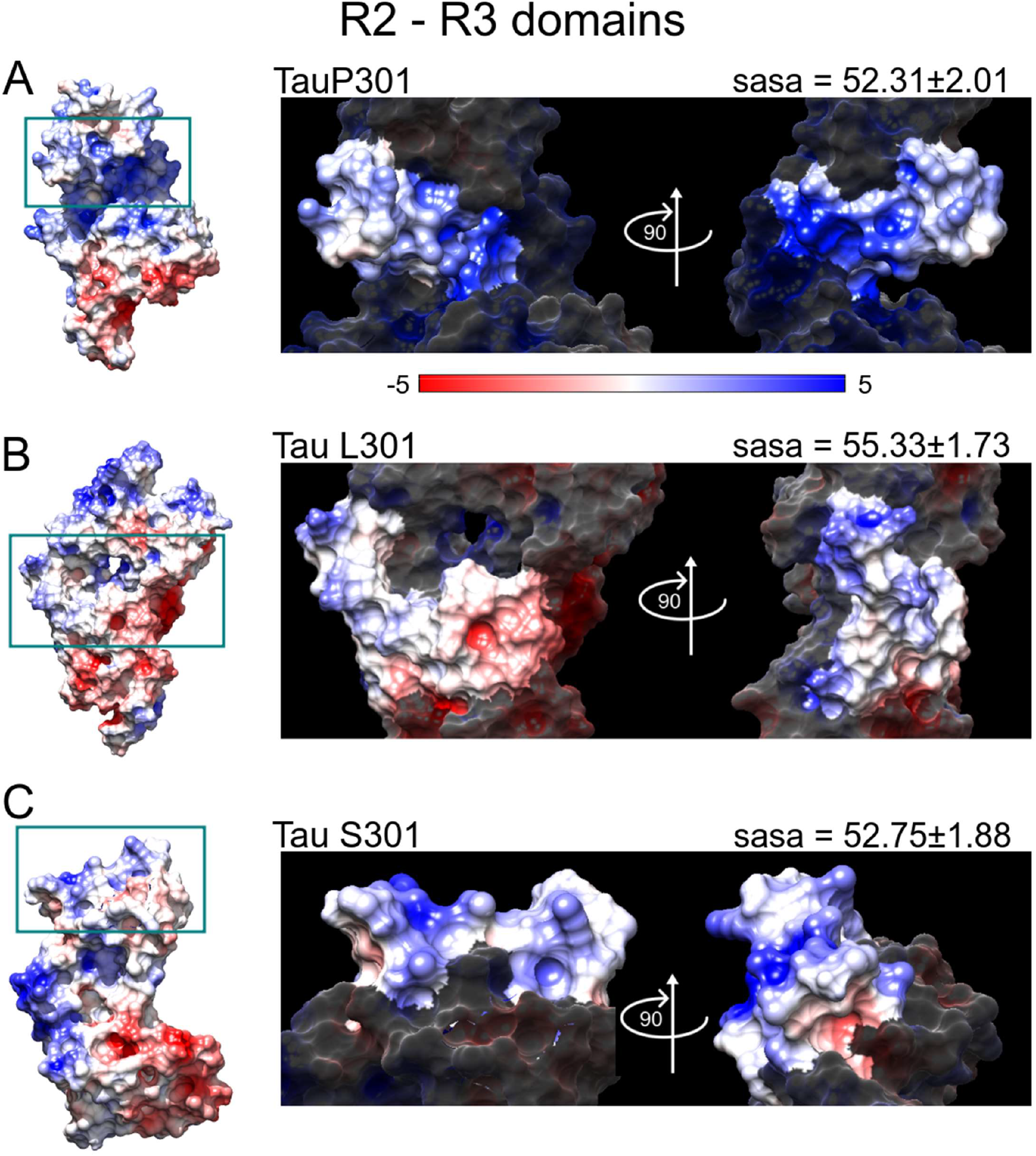
Energy analysis of the R2-R3 domains. **(A)** RMSD plot of Tau 3D structures obtained after 500 ns of MD simulations. **(B)** Final alignment of the R2-R3 domains using the 3D structure of the Tau wild-type (orange), Tau^P301S^ (green) and Tau^P301L^ (blue). **(C)** Radius of gyration plot shows the folding of the Tau 3D proteins **(D)** Construction of the free energy landscape maps (FELs) using the data obtained from the RMSD and radius of gyration as variables. The figure only shows the results obtained for the R2 and R3 domains from the last 100ns of simulation. **(E)** Energy interactions of R2-R3 domains.

**Figure S5.**
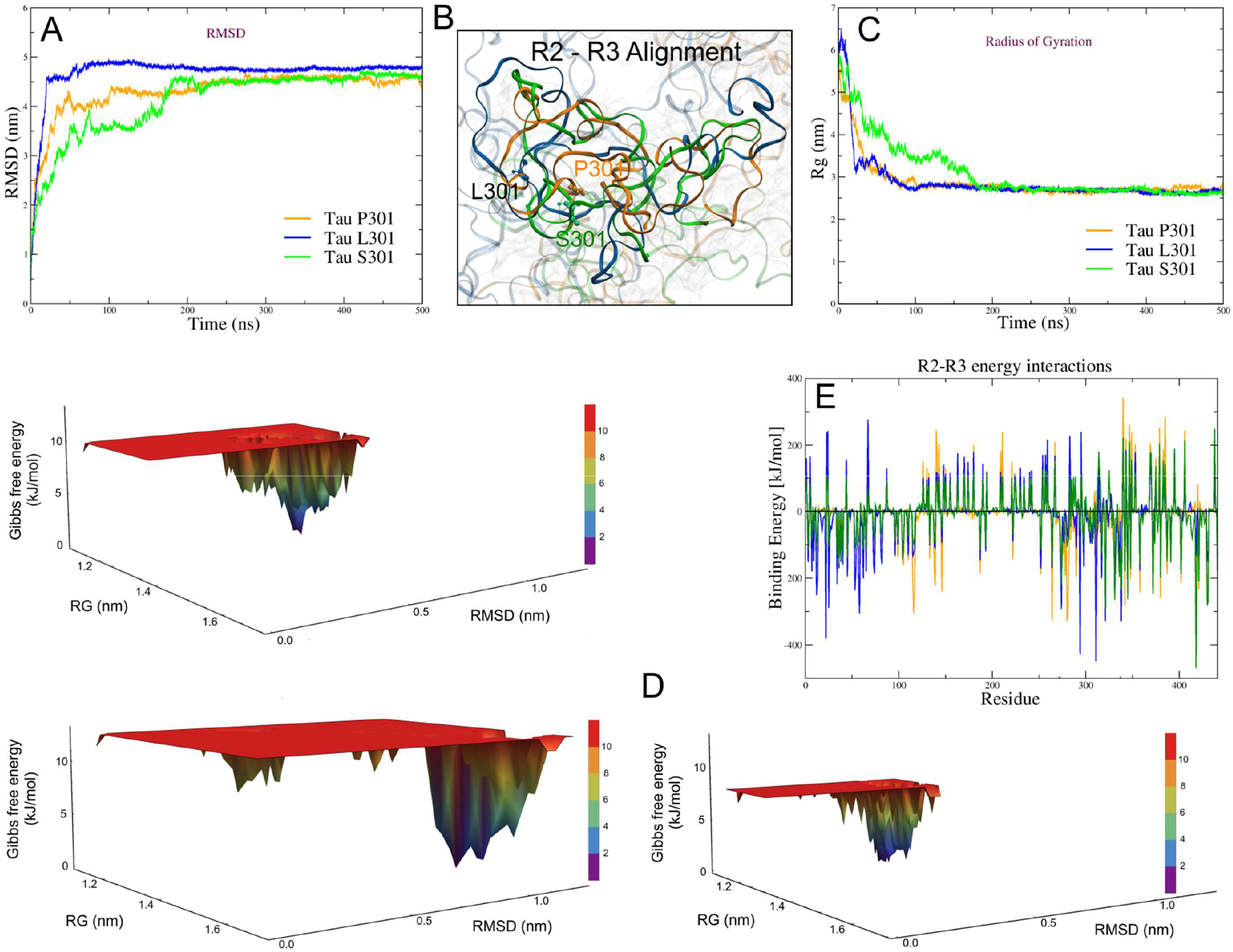
S262 site molecular analysis. **(A)** Energetic molecular interactions and SASA on the Tau protein surfaces. The Tau^P301L^ protein shows the greatest solvent accessible area and bigger destabilization energy compared to other Tau proteins. **(B)** Energy contribution per residue in the interactions with the S262 residue. The blue box indicates interaction regions.

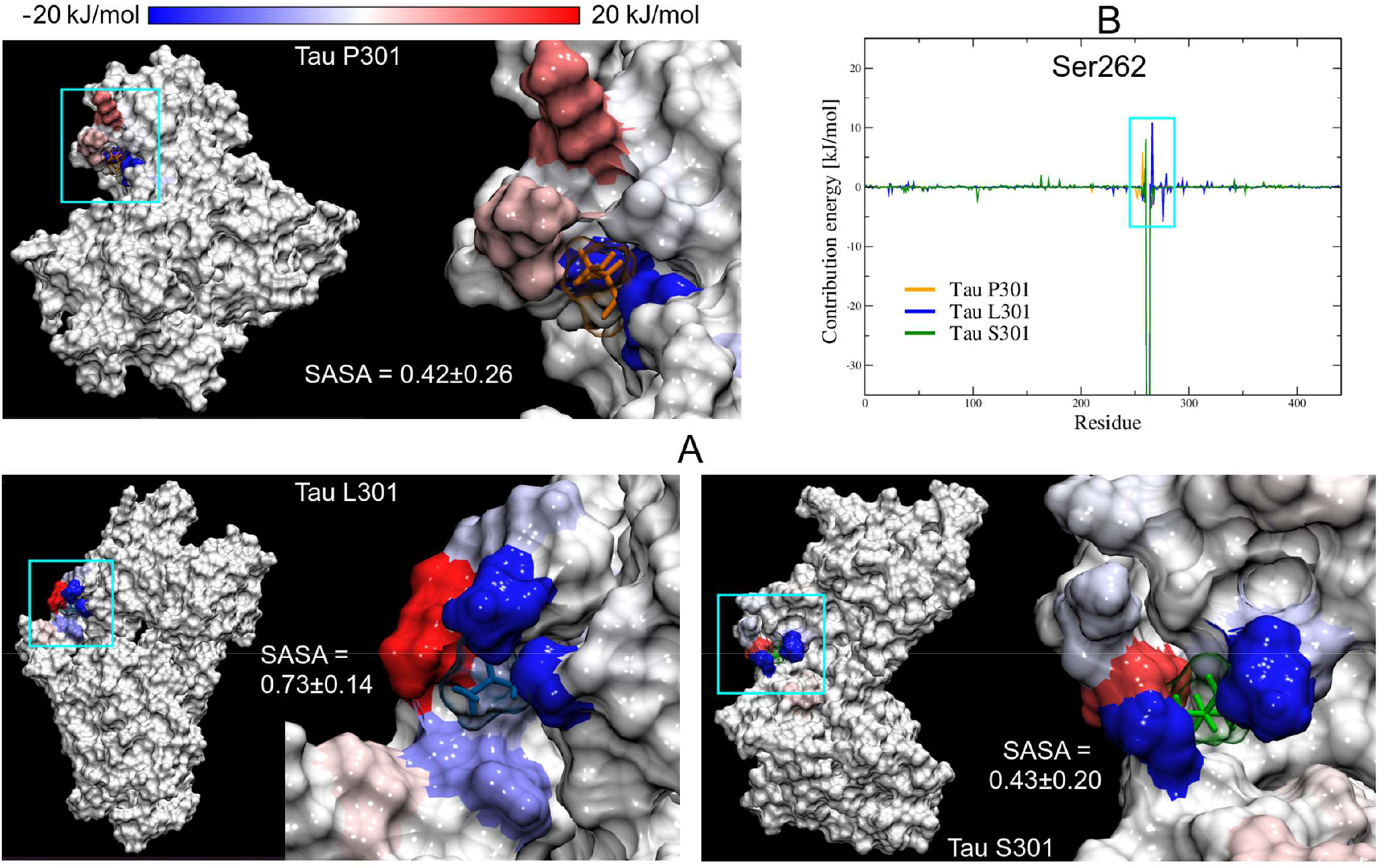

## SUPPLEMENTARY MATERIAL

**Table S1.**

System Details
| System | Total atoms | Protein atoms | Solvent molecules | Box size |
| --- | --- | --- | --- | --- |
| Tau P301 | 300427 | 6416 | 98003 | 16.98 x 15.14 x 11.71 |
| Tau L301 | 294825 | 6421 | 96134 | 16.74 x 15.47 x 11.39 |
| Tau S301 | 318859 | 6413 | 104148 | 17.11 x 15.84 x 11.77 |

Stability Descriptors
| System | RMSD | RMSF | Radius of Gyration | H Bonds |  |
| --- | --- | --- | --- | --- | --- |
|  |  |  |  | Intra | Inter |
| Tau P301 | 4.23±0.64 | 1.30±0.45 | 2.95±0.60 | 209±21 | 1142±53 |
| Tau L301 | 4.58±0.74 | 1.29±0.40 | 2.99±0.83 | 228±26 | 1098±70 |
| Tau S301 | 4.12±0.70 | 1.16±0.50 | 3.10±0.69 | 218±15 | 1124±41 |

**Table S2.** Surface areas.

Solvent accessible surface areas (nm<sup>2</sup>)
| Tau Protein |  | Total area | R2 - R3 | Tyr18 | Thr181 | Ser202 | Thr 205 | Ser262 | X301 | Ser356 |
| --- | --- | --- | --- | --- | --- | --- | --- | --- | --- | --- |
| P301 | final val. | 269.79±6.10 | 52.31±2.01 | 0.87±0.25 | 0.31±0.18 | 0.56±0.16 | 1.09±0.29 | 0.42±0.26 | 0.55±0.13 | 0.96±0.11 |
|  | Initial val. | 360.99 | 46.05 | 0.22 | 1.00 | 1.16 | 0.31 | 0.82 | 0.29 | 0.93 |
| L301 | final val. | 240.31±4.65 | 55.33±1.73 | 0.28±0.20 | 0.26±0.10 | 0.17±0.13 | 0.55±0.10 | 0.73±0.14 | 0.11±0.06 | 0.04±0.03 |
|  | Initial val. | 361.59 | 51.94 | 1.72 | 1.09 | 0.77 | 0.16 | 0.84 | 0.78 | 0.24 |
| S301 | final val. | 250.35±5.80 | 52.75±1.88 | 0.22±0.17 | 0.12±0.06 | 0.23±0.14 | 0.42±0.11 | 0.43±0.20 | 0.13±0.05 | 0.23±0.11 |
|  | Initial val. | 362.33 | 52.05 | 1.54 | 1.07 | 0.95 | 0.17 | 0.88 | 0.31 | 0.26 |

## DATA SOURCES

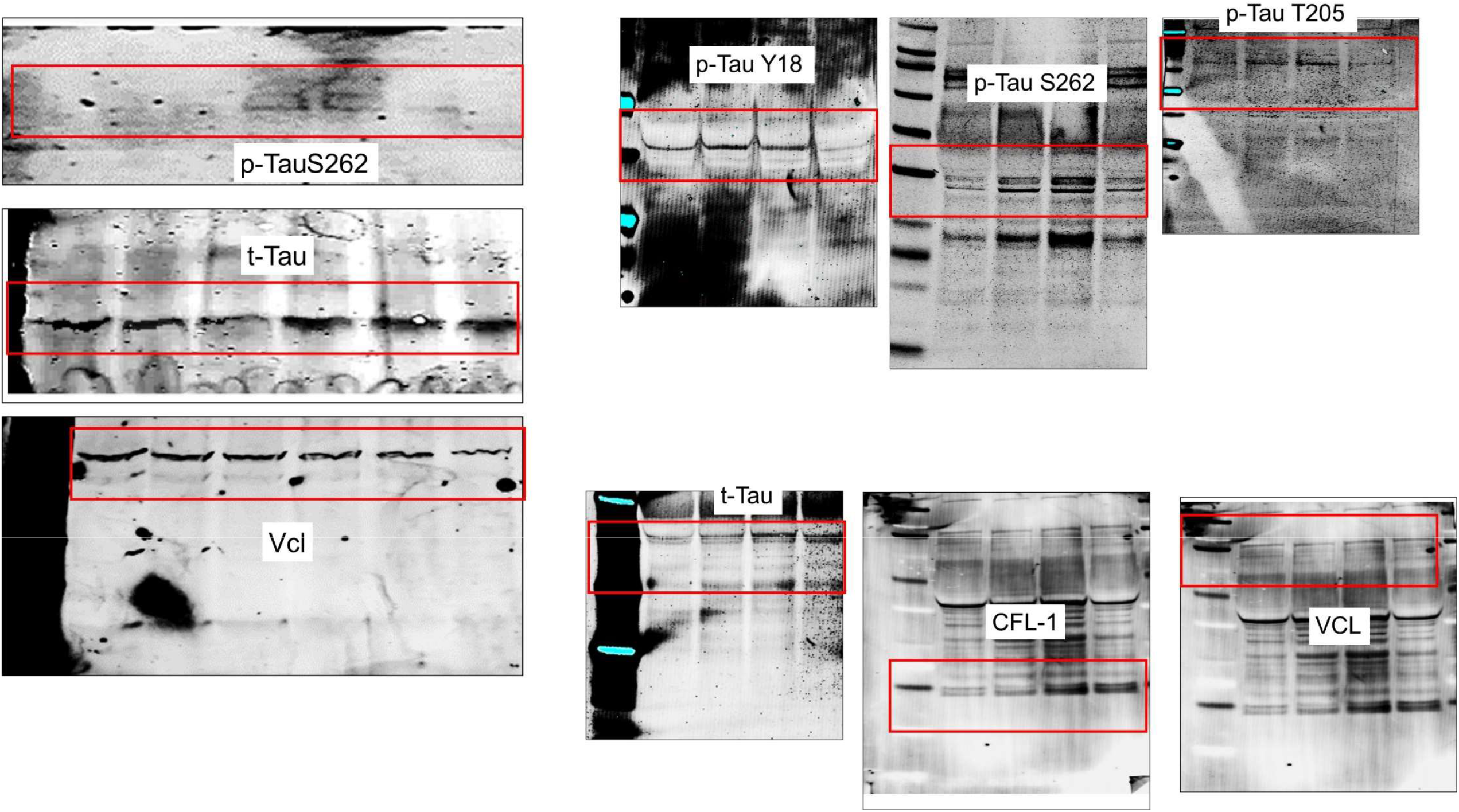

## Notes

### Competing Interest Statement

The authors have declared no competing interest.

